# FARL-11 (STRIP1/2) is Required for Sarcomere and Sarcoplasmic Reticulum Organization in *C. elegans*

**DOI:** 10.1101/2023.03.05.531173

**Authors:** Sterling C.T. Martin, Hiroshi Qadota, Andres F. Oberhauser, Jeff Hardin, Guy M. Benian

## Abstract

Protein phosphatase 2A (PP2A) functions in a variety of cellular contexts. PP2A can assemble into four different complexes based on the inclusion of different regulatory or targeting subunits. The B’’’ regulatory subunit “striatin” forms the STRIPAK complex consisting of striatin, a catalytic subunit (PP2AC), striatin interacting protein 1 (STRIP1), and MOB family member 4 (MOB4). In yeast and *C. elegans,* STRIP1 is required for formation of the endoplasmic reticulum (ER). Since the sarcoplasmic reticulum (SR) is the highly organized muscle-specific version of ER, we sought to determine the function of the STRIPAK complex in muscle using *C. elegans*. CASH-1 (striatin) and FARL-11 (STRIP1/2) form a complex *in vivo*, and each protein is localized to SR. Missense mutations and single amino acid losses in *farl-11* and *cash-1* each result in similar sarcomere disorganization. A missense mutation in *farl-11* shows no detectable FARL-11 protein by immunoblot, disruption of SR organization around M-lines, and altered levels of the SR Ca^+2^ release channel UNC-68.

**Summary:** Protein phosphatase 2A forms a STRIPAK complex when it includes the targeting B’’’ subunit “striatin” and STRIP1. STRIP1 is required for formation of ER. We show that in muscle STRIP1 is required for organization of SR and sarcomeres.

## Introduction

A major serine-threonine protein phosphatase that is conserved from yeast to humans is protein phosphatase 2A (PP2A; Shi, 2009). PP2A functions in a wide variety of cellular contexts and forms multiple specific protein complexes via different regulatory or targeting subunits. The PP2A core enzyme comprises a catalytic subunit C (PP2AC) and a scaffolding subunit A (PP2AA). This core forms a complex with a variety of regulatory B subunits, with the B subunits mediating subcellular localization and/or substrate recognition. There are four families of regulatory B subunits: B (B55 or PPP2R2), B’ (B56 or PPP2R5), B” (B72 or PPP2R3), and B’’’ (striatins).

The striatin-interacting phosphatase and kinase (STRIPAK) complex is a PP2A complex in which the regulatory subunit is the protein striatin. Recently, the structure of the STRIPAK complex was solved at 3.2 Å resolution using cryo-EM (Jeong et al., 2021). The complex consists of a catalytic subunit (PP2AC), a scaffolding subunit (PP2AA) called “striatin”, striatin interacting protein 1 (STRIP1), and MOB family member 4 (MOB4). The striatin coiled-coil domain forms an elongated scaffold that links the complex together.

In the budding yeast, *Saccharomyces cerevisiae*, the “Far” complex, originally identified as crucial for pheromone-induced cell cycle arrest, is the yeast counterpart of the STRIPAK complex. The Far complex localizes to the endoplasmic reticulum (ER), and it has been shown that the six Far subunits (Far3, 7, 8, 9, 10 and 11) assemble at the ER in a specific sequence (Pracheil and Liu, 2013). Similarly, in *C. elegans*, the nematode ortholog of yeast Far11, called FARL-11, is localized to the ER and outer nuclear membrane of early embryos and the germline (Maheshwari et al., 2016). Moreover, in a *farl-11* loss of function mutant, the morphology of the ER is severely disrupted, indicating that FARL-11 protein is required for normal ER morphology. In striated muscle, the sarcoplasmic reticulum (SR) is a highly organized, muscle-specific version of the ER, and is the storage and release depot for Ca^2+.^ In response to an action potential from motor neurons, the calcium release channel of the SR (the ryanodine receptor) is activated and opens to allow flow of Ca^2+^ from the SR into the muscle cell cytoplasm where Ca^2+^ interacts with components of the sarcomere (usually the troponin/tropomyosin complex) to activate muscle contraction.

The role of STRIPAK in SR function has not been determined. We turned to *C. elegans* as a genetically tractable model organism in which to investigate the function of STRIPAK in muscle function. *C. elegans* is an outstanding system for discovering new conserved aspects of muscle assembly, maintenance and regulation (Gieseler et al., 2017). In a previous study we reported that loss of function of most single components of PP2A results in sarcomere disorganization (Qadota et al., 2018). This includes the catalytic subunit C (LET-92), the scaffolding subunit A (PAA-1), regulatory subunit B (SUR-6), regulatory subunit B’ (either PPTR-1 or PPTR-2), regulatory subunit B” (RSA-1) and regulatory subunit B’’’ (CASH-1). Moreover, we reported that with available antibodies to five of these PP2A components, we were able to localize the components to different and overlapping components of the sarcomere. For example, SUR-6 and PPTR-1 localize to I-bands, PPTR-2 localizes to M-lines and dense bodies (Z-disks), and RSA-1 localizes to M-lines and I-bands (Qadota et al., 2018).

Here, we used CRISPR-tagging, antibodies and genetic mutants to investigate the role of the STRIPAK complex in adult *C. elegans* muscle. We report that FARL-11 (STRIP1/2) and CASH-1 (STRN3) are localized to the SR, and that the phenotypes of *farl-11* and *cash-1* loss of function mutants are similar. Either mutant has substantial effects on sarcomere organization, and at least a *farl-11* mutant disrupts SR organization, especially around M-lines.

## Results

### *Mutations in* cash-1 (STRN3) *and* farl-11 (STRIP1/2) *cause sarcomeric organizational defects*

To facilitate our studies, we generated antibodies to a C-terminal portion of FARL-11, and also used CRISPR/Cas9 to generate an N-terminal fusion of FARL-11 to mNeonGreen and Flag (strain SU853), and an N-terminal fusion of CASH-1 to TagRFP and myc (strain SU854; Figure 1A). As shown in Figure 1B, anti-FARL-11 rabbit polyclonal antibodies react to a protein of the expected size (∼110 kDa) from wild-type, and an appropriately larger fusion protein from the mNeonGreen-tagged strain. Antibodies to myc detect an appropriately sized protein (∼130 kDa) from the strain that expresses an RFP- and myc-tagged fusion to CASH-1. To seek further evidence of the evolutionary conservation of the STRIPAK complex, we asked whether an *in vivo* complex could be detected in nematodes that contains both CASH-1 (STRN3) and FARL-11 (STRIP1/2). As shown in Figure 1C, immunoprecipitation of TagRFP-myc-CASH-1 from an extract of strain SU854, co-immunoprecipitates FARL-11.

**Figure 1.**
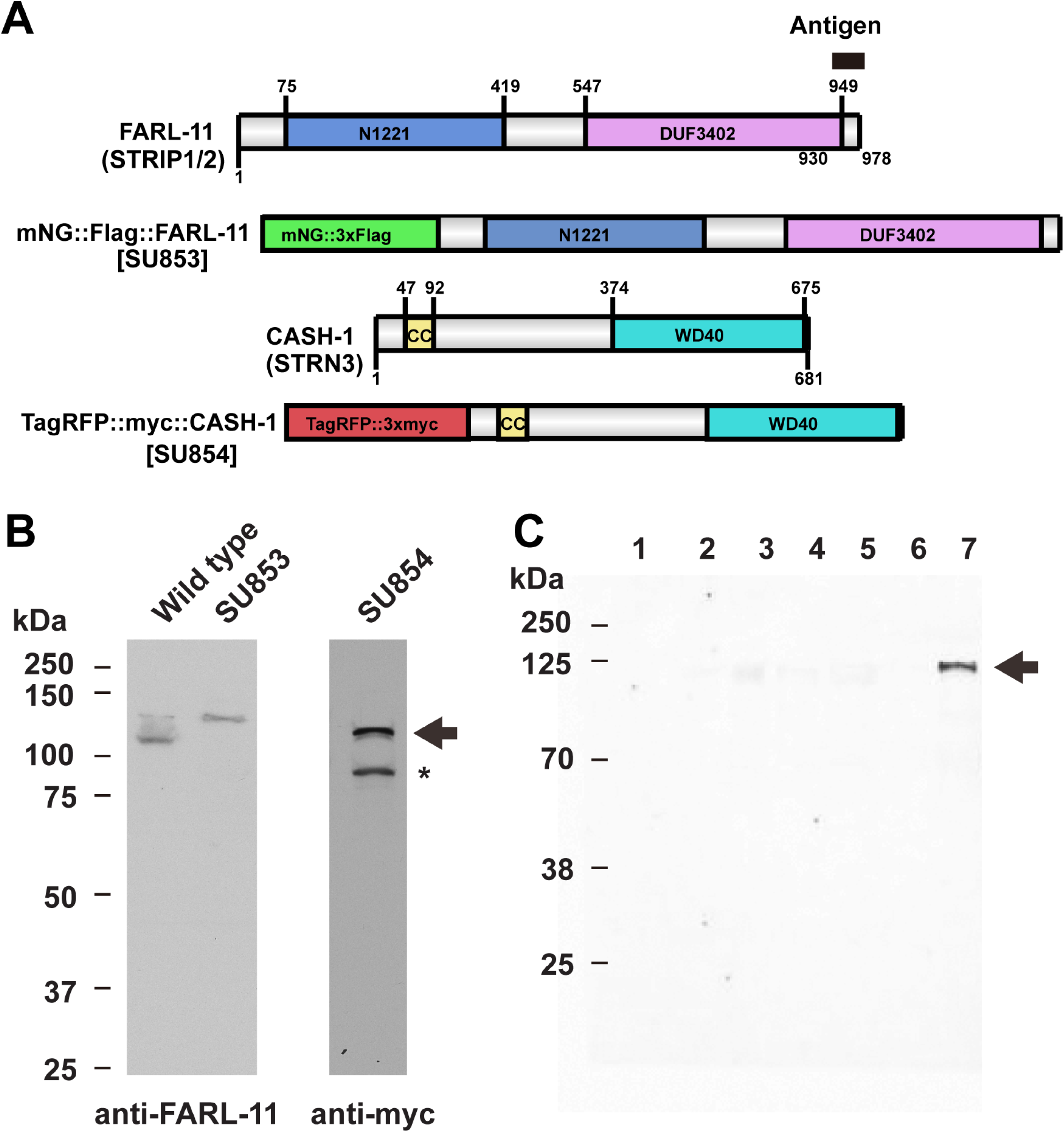
Domain organization, tagging design, detection of expression of FARL-11 and CASH-1, and coIP of FARL-11 with CASH-1. (A) Domain organization of FARL-11 and CASH-1 and location of tags. CRISPR/Cas9 was used to create nematode strain SU853, in which consecutive mNeonGreen and Flag tags were added to the N-terminus of FARL-11, and strain SU854, in which RFP and myc tags were added to the N-terminus of CASH-1. “Antigen” refers to the 50 amino acid peptide that was used to generate rabbit antibodies to FARL-11. (B) Western blots demonstrating detection of FARL-11, mNG-Flag-FARL-11, and RFP-myc-CASH-1. Nematode extracts were run on a gel, transferred to membrane and reacted against the indicated antibodies. The major bands detected using anti-FARL-11 run at sizes expected for endogenous and tagged FARL-11. Using anti-myc, two bands are detected from SU854. The band indicated with the arrow runs at the size expected for RFP-myc-CASH-1. The band indicated with an asterisk is a protein of nematode or bacterial origin that also reacts with antibodies to myc. (C) Co-immunoprecipitation of FARL-11 with CASH-1. As described in materials and methods, TagRFP-myc-CASH-1 was immunoprecipitated from a lysate of strain SU854, and proteins from consecutive steps of the procedure were separated by SDS-PAGE, blotted and incubated with anti-FARL-11. Lanes: (1) total lysate, (2) insoluble proteins pelleted from total lysate, (3) clarified cell lysate, (4) proteins associated with magnetic agarose control beads, (5) proteins that did not bind to anti-RFP magnetic agarose, (6) proteins in wash of anti-RFP magnetic agarose, (7) proteins eluted from anti-RFP magnetic agarose. Arrow indicates the position of FARL-11. For parts (B) and (C), the positions of molecular weight size markers are indicated.

In our previous report of the PP2A complex in *C. elegans* muscle, we showed that RNAi knockdown of CASH-1 (STRN3) resulted in adults that are slow-moving and sterile, and by immunostaining with antibodies to myosin, these animals have disorganized sarcomeric A-bands (Qadota et al. 2018). Inspection of the Million Mutation Project (MMP) collection of mutant strains (Thompson et al., 2013) revealed several strains that have missense mutations in conserved residues of CASH-1, but one of them, *cash-1(gk705529)*, which results in a H121Y mutation, had severe phenotypes: a high percentage of embryonic lethality, and, among the animals that reached adulthood, greatly reduced motility. To avoid the confounding influence of the many background mutations in a MMP strain, we re-created the H121Y mutation by CRISPR in our RFP- and myc-tagged CASH-1 strain (Figure 2A). The crystal structure of human striatin 3 coiled-coil domain revealed that it forms a parallel dimer and that two highly conserved residues, L121 and L125, are critical for homodimerization and binding to the A subunit of PP2A (Chen et al., 2014). We used CRISPR to mutate the equivalent leucines (L82 and L86) to glutamates in our RFP- and myc-tagged CASH-1 strain (Figure 2A). Strains containing the H121Y mutation were adult viable and fertile. However, nematodes homozygous for the L82E / L86E mutations reached adulthood but were sterile, and thus we maintained the mutant chromosome over a balancer chromosome. We immunostained adults from each strain with antibodies to UNC-89 (obscurin) to assess M-lines, myosin heavy chain A (MHC A) to assess A-bands, and UNC-95 to assess the bases of the M-lines and dense bodies (Z-disks) and muscle cell boundaries. As shown in Figure 2B, each of the mutants display disorganization of these structures, with a more severe phenotype in the L82E/L86E mutant strain. It is interesting that for the L82E / L86E strain some regions of muscle cells show unusually broad localization of MHC A and especially UNC-89 (indicated by arrows). In addition, UNC-95 staining is moderately disorganized in each mutant strain, especially with respect to the dense bodies and muscle cell boundaries. To assess to what extent the sarcomeric defects could be attributed to a decreased level of mutant protein vs. functionally defective proteins, we compared the level of the RFP/myc-CASH-1 in an otherwise wild-type background to the level of RFP/myc-CASH-1 H121Y, and RFP/myc-CASH-1 L82E/L86E. As shown in Figure 2C and D, the levels of RFP/myc-CASH-1 were similar in wild type as compared to either H121Y or L82E/L86E mutants. Thus, the sarcomeric disorganization is most likely due to decreased function in CASH-1 due to these missense mutations.

**Figure 2.**
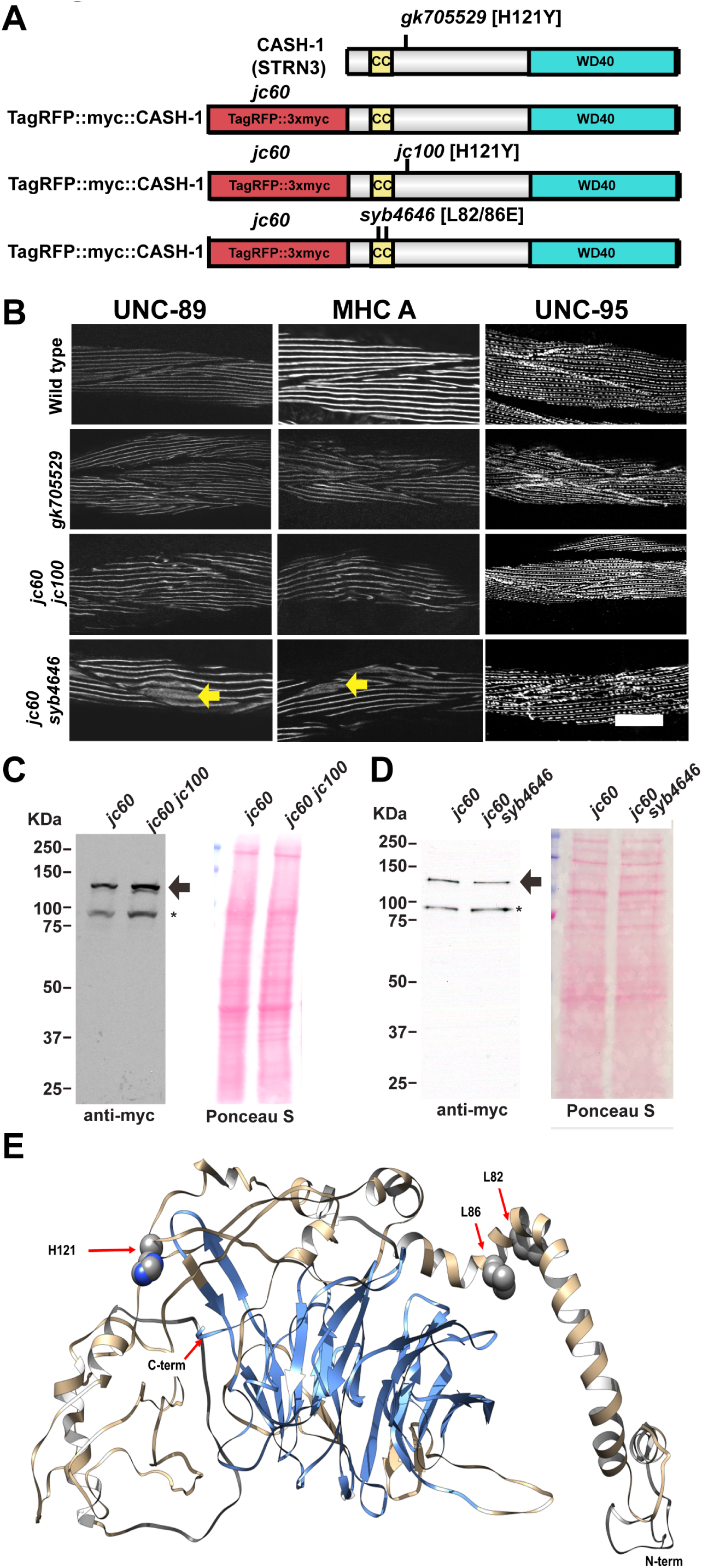
Missense mutations in *cash-1* result in mild to moderate disorganization of muscle sarcomeres. (A) Schematic representation of domains in CASH-1 and tagged CASH-1. The H121Y mutation was originally found in the MMP allele *cash-1(gk705529)* and re-created by CRISPR/Cas9 in the RFP-myc tagged *cash-1* strain *jc60*, as *jc60 jc100*. The double missense mutant L82E / L86E was also created by CRISPR/Cas9 in the RFP-myc taged *cash-1* strain *jc60*, as *syb4646*. (B) Representative images of several body wall muscle cells from wild type, *cash-1(gk705529)*, *cash-1(jc60 jc100)*, and *cash-1(syb4646)*, immunostained with antibodies to UNC-89 for M-lines, MHC A myosin for A-bands, and UNC-95 for bases of M-lines, dense bodies and muscle cell boundaries. Mild disorganization of these structures is seen in both strains containing the H121Y missense mutation, with the more severe effect seen with UNC-95 staining, especially at the muscle cell boundaries. A more severe effect on sarcomere and IAC organization is seen in the L82E/L86E double mutant. Yellow arrows point out widening of the localization of UNC-89 and MHC A. Scale bar, 10 μm. (C) and (D) Western blot analysis of the level of RFP-myc-CASH-1 in *cash-1(jc60)* and *cash-1(jc60 jc100)* in C, and *cash-1(syb4646)* in D. For each, on the left, reaction with anti-myc; on the right, the Ponceau S staining of the blot. Arrow indicates the band expected from RFP-myc-CASH-1; asterisk denotes a cross-reacting band. The presence of the H121Y mutation or the L82E / L86E double mutations do not seem to affect the level of steady state RFP-myc-CASH-1. (E) Structure of CASH-1 based on the human striatin 3 3D structure (PDB: 7K36) (Jeong et al., 2021) modelled with SWISS-MODEL (Waterhouse et al., 2018), and showing residues L82, L86 and H121. The colors represent different putative WD40 repeats (in blue) and disordered region (in grey).

To obtain further insight into how these missense mutations affect the function of the CASH-1 protein, we generated a homology model of CASH-1 based on the cryoEM structure of the orthologous human protein, STRN3 (Jeong et al. 2021). Figure 2F displays the locations of the affected residues on this homology model. In order to predict the potential effects of the mutations we used the rotamer/mutation tool in Chimera (Pettersen *et al*., 2004) and then energy minimized to visualize interatomic clashes and contacts based on van der Waals radii. In addition, we used the DynaMut2 (Rodrigues *et al.,* 2021) to predict the effects of the point mutations on protein stability and dynamics. L82 and L86 lie along the surface of a long α-helix (as expected), and we predict that the L82E and L86E mutations will have no significant effect on structure. Nevertheless, as shown for the STRN3, these mutations are likely to affect homodimerization. H121 resides in a loop connecting two α-helices. Given that H121 lies on the surface of the protein, the H121Y could conceivably affect binding to FARL-11.

We selected one *farl-11* MMP allele for analysis, *farl-11(gk437008)*, which has a non-conservative L292 to P change in the N1221 domain. This strain was outcrossed 3X to wildtype. Although we could not create this mutation using CRISPR, we did generate two additional mutations from the CRISPR procedure at nearby residues in this same domain: *farl-11(jc93)*, which deletes the single amino acid T281 in-frame, and *farl-11(jc94)*, which deletes the single amino acid P280 in-frame. After outcrossing 2X to wildtype and immunostaining, all three mutant alleles show severely disorganized sarcomeres (Figure 3B). Of particular note is the wider distribution of both UNC-89 and myosin, similar to what we observed in the *cash-1* mutant containing the L82E / L86E mutations. For the *farl-11* mutants, this wider distribution of UNC-89 and MHC A can be discerned more clearly in the zoomed insets (Figure 3C). To assess to what extent the sarcomeric defect is due to a decreased level of mutant protein vs. a functionally defective mutant protein, we compared the levels of FARL-11 protein from all three *farl-11* mutants to wildtype. As shown in Figure 3D, Western blot analysis indicates no detectable FARL-11 protein in *farl-11(gk437008)* (L292P). Moreover, both *jc93* and *jc94* show decreased levels of FARL-11: *jc93* has 48.9 % of the wild-type level, and *jc94* has 64.1% of the wild-type level (mean, n = 4). We were somewhat surprised that *gk437008*, a missense mutant, shows no detectable FARL-11 protein. There are no other mutations in the *farl-11* gene in this strain (VC30204), based on a query of the MMP website at Simon Fraser University (http://genome.sfu.ca/mmp/).

**Figure 3.**
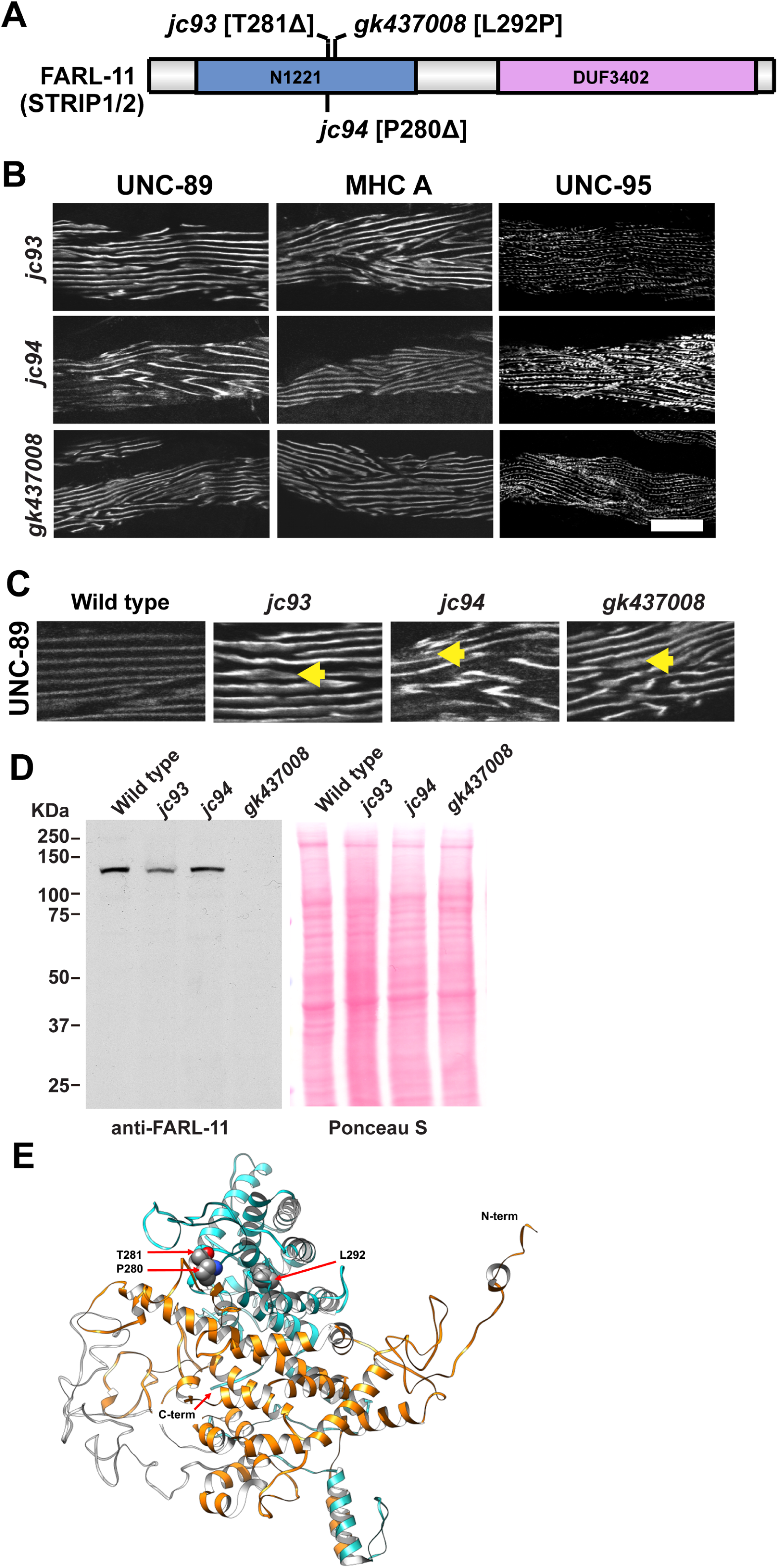
Missense mutations in *farl-11* result in disorganization of sarcomeres. (A) Schematic representation of domain organization of FARL-11 and location of mutations. (B) Representative images of several body wall muscle cells from each of *farl-11(jc93)*, *farl-11(jc94)*, and *farl-11(gk437008)*, immunostained with antibodies to UNC-89 for M-lines, MHC A myosin for A-bands, and UNC-95 for bases of M-lines, dense bodies and muscle cell boundaries. As compared with wild type (top row, Figure 2B), organization of all of these structures is affected. Scale bar, 10 μm. (C) 2X zoomed-in views of immunostaining with antiUNC-89 showing the moderate to severe mislocalization of UNC-89 in all three mutants compared to wild type. Arrows indicate the abnormal broadening of localization. (D) Western blot analysis of the level of FARL-11 in wild type as compared to *farl-11* mutants. On the left, reaction with anti-FARL-11; on the right, the Ponceau S staining of the blot. Although deletion of single amino acids in each of the mutants *jc93* and *jc94* does not significantly affect the level of the mutant FARL-11 protein, the L292P substitution in *gk437008* results in no detectable mutant FARL-11 protein. (E) Structure of FARL-11 based on the human STRIP1 3D structure (PDB: 7K36) (Jeong et al, 2021) modelled with SWISS-MODEL (Waterhouse et al., 2018), and showing residues T281, P280 and L292. The colors represent different putative domains (N1221 in cyan and DUF3402 in orange) and disordered region (in grey).

To obtain additional understanding about how these mutations might affect the function of FARL-11, we generated an homology model of FARL-11 based on the cryoEM structure of the human ortholog, STRIP1 (Jeong et al. 2021). Figure 3E depicts the locations of the affected residues in this structure. P280 and T281 are located in a highly disordered 47 amino acid loop, and L292 is located in the center of an α-helix. Like for CASH-1 we used the rotamer/mutation tool in Chimera (Pettersen *et al*., 2004) and the DynaMut2 web tool (Rodrigues et al., 2021) to predict the impact of the mutations on protein stability.

We predict that deletion of either P280 or T281 will have minimal effect on the structure or stability of the protein. This prediction is consistent with the approximately normal abundance of these mutant proteins lacking either residue observed by western blot (Figure 3D). In contrast, the L292P mutation leads to several interatomic clashes and hence is likely to break the α-helix and lead to a local and even global change in the structure of the protein and lower its stability (ΔΔG^Stability^ = −0.72 kcal/mol). Again, our western blot result on this L292P mutant is consistent with the prediction in that we cannot detect any FARL-11 L292P protein in worm lysates (Figure 3D).

### CASH-1 and FARL-11 localize near dense bodies

We next sought to determine the localization of CASH-1 and FARL-11 in nematode body wall muscle. To localize CASH-1 we utilized our CRISPR strain, SU854, which expresses an RFP/myc-CASH-1 fusion protein. As shown in Figure 4A, in which CASH-1 was detected with antibodies to myc and M-lines and dense bodies detected with antibodies to UNC-95, CASH-1 is localized between and surrounding dense bodies. To localize FARL-11 we used antibodies to FARL-11. As indicated in Figure 4B, in which we co-stained with antibodies to PAT-6 (α-parvin), which also localizes to the bases of M-lines and dense bodies, FARL-11 localizes between dense bodies and also surrounds dense bodies, but not as broadly as CASH-1. Unfortunately, we were not able to address whether CASH-1 and FARL-11 co-localize, despite several attempts. This is probably because the anti-myc-CASH-1 staining produces a low signal and high background.

**Figure 4.**
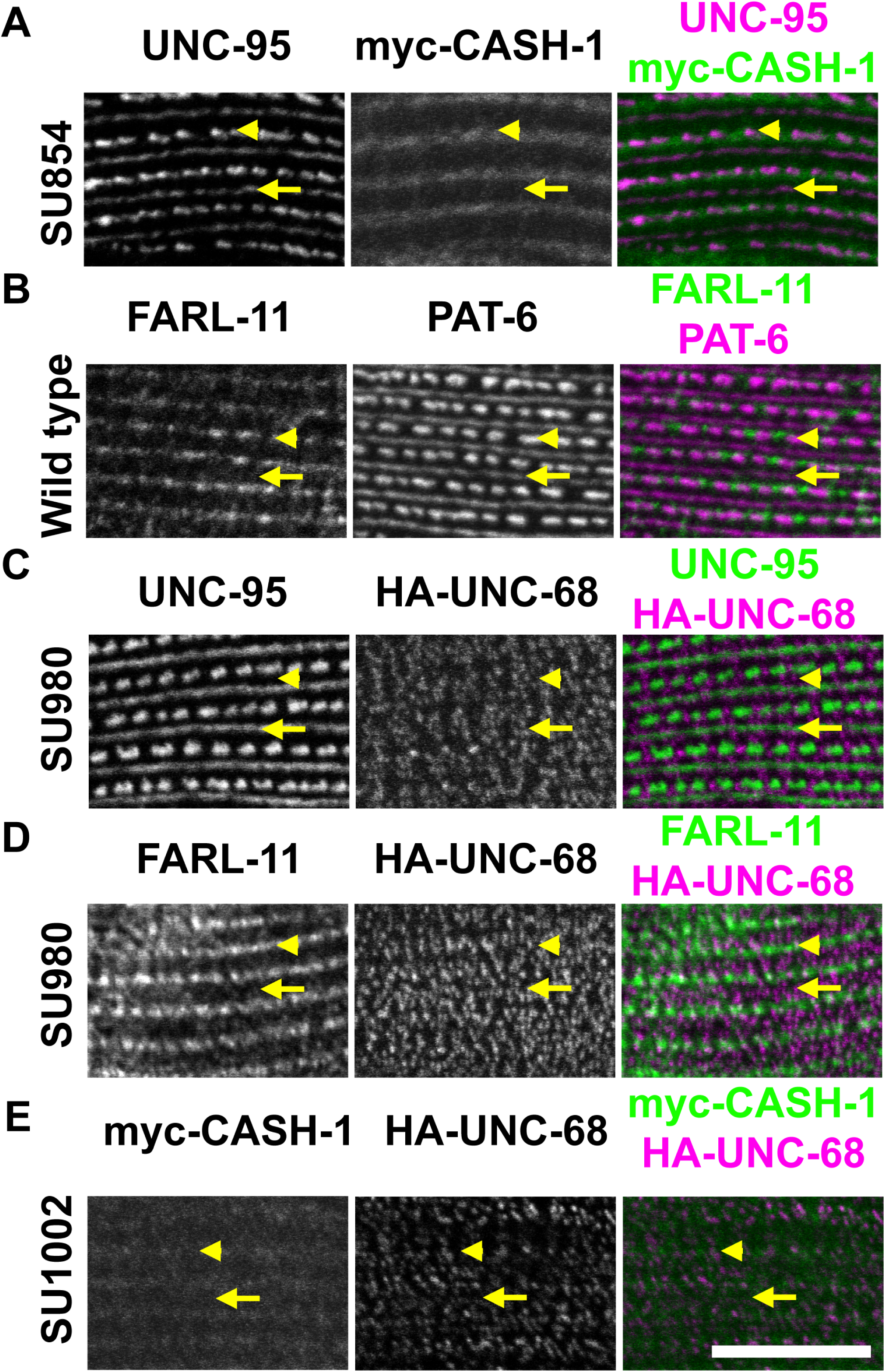
Localization of CASH-1, FARL-11 and UNC-68 by confocal microscopy. Each panel shows a portion of a single body wall muscle cell immunostained with antibodies to the indicated proteins or to the myc or HA tags. Yellow arrow heads point to a row of dense bodies; yellow arrows to single M-lines. Scale bar, 10 μm. (A) Strain SU854 (expresses RFP-myc-CASH-1) showing localization of UNC-95 and myc-CASH-1. Myc-CASH-1 localizes between and surrounding dense bodies. (B) Wild type showing localization of PAT-6 and FARL-11. FARL-11 also localizes between and surrounding dense bodies. (C) Strain SU980 (expresses HA-UNC-68) showing localization of UNC-95 and HA-UNC-68. HA-UNC-68 localizes to puncta organized in a striated pattern roughly surrounding dense bodies and M-lines. (D) Strain SU980 showing localization of FARL-11 and HA-UNC-68. The two proteins show partial co-localization suggested by the white puncta created by overlap of green FARL-11 and magenta HA-UNC-68. (E) Strain SU1002 (expresses myc-CASH-1 and HA-UNC-68) showing localization of myc-CASH-1 and HA-UNC-68. The two proteins partially co-localize. Scale bar, 10 μm.

### FARL-11 localizes to the sarcoplasmic reticulum

Transmission electron micrographs have revealed that the SR in *C. elegans* body wall muscle is restricted to thin membranous sacs closely adjacent to the dense bodies as well as near the bases of M-lines or the middle of the A-band region (Gieseler et al. 2017). In *C. elegans*, the ryanodine receptor or calcium release channel of the SR is encoded by a single gene, *unc-68* (Maryon et al., 1996; Sakube et al., 1997). Using confocal microscopy to image localization of antibodies to UNC-68 and various components of the myofilament lattice (MHC A, MHC B, vinculin, α-actinin), Maryon et al. (1998) concluded that UNC-68 resides primarily in flattened vesicular sacs adjacent to the outer muscle cell membrane in the A-band region. In contrast, Hamada et al. (2002), using antibodies to UNC-68 and either rhodamine-phalloidin or antibodies to the A-band protein paramyosin, concluded that UNC-68 resides in the I-band region. Although these two studies seem to give different localizations for UNC-68, together, the results include both known or suspected locations of the SR (surrounding dense bodies and adjacent to base of M-lines). Unfortunately, the antibodies generated to UNC-68 utilized in these studies are no longer available. A recent report by Piggott et al. (2021) provides the best confocal images of UNC-68 localization in muscle. These authors created a split-GFP knock-in allele for *unc-68* and their confocal images show that GFP::UNC-68 localizes to rows of puncta, some large, some small, but the authors did not co-localize with any sarcomeric markers.

Given that FARL-11 has been localized to the ER of early nematode embryos and that the SR is a muscle-specific type of ER, we used CRISPR/Cas9 to create strain SU980, in which UNC-68 is tagged at its N-terminus with 3xHA. By conventional confocal microscopy, HA-UNC-68 exists in puncta which are organized in a repeating striated pattern roughly surrounding both dense bodies and M-lines (Figure 4C). We next asked whether FARL-11 and UNC-68 might co-localize. As shown in Figure 4D, there is at least some co-localization of the two proteins, suggested by the white puncta created by the overlap of green FARL-11 and magenta HA-UNC-68 signals. Similarly, HA-UNC-68 and myc-CASH-1 partially co-localize (Figure 4E).

To obtain more information about the localization of UNC-68, we used structured illumination microscopy (SIM), which has an ∼120 nm resolution in the XY plane. SIM followed by 3D reconstruction (Figure 5A) shows that HA-UNC-68 localizes to a series of puncta, very similar to the images reported by Piggot et al. (2021). Co-staining with PAT-6 (α-parvin), which localizes to the bases of the M-lines and dense bodies reveals that the HA-UNC-68 puncta localize on either side of the M-lines, and surround the dense bodies, with the larger puncta being closer to the dense bodies. There is also accumulation of HA-UNC-68 at muscle cell boundaries (indicated by yellow arrows in Figure 5B). When a 3D reconstruction of an entire muscle cell is viewed on its side, we observe that in addition to being localized like PAT-6 close to the muscle cell membrane, there is significant localization of HA-UNC-68 in the deeper portions of the muscle cell (Figure 5C). To determine how deeply this HA-UNC-68 is located, we co-stained anti-HA with anti-ATN-1 or with anti-UNC-89. ATN-1 localizes to the deepest portions of the dense bodies and defines the deepest portion of the myofilament lattice. As can be seen in Figure 5D, HA-UNC-68 is found mainly near the outer muscle cell membrane but also extends more deeply into the muscle cell than the myofilament lattice. A similar result was observed when HA-UNC-68 was co-stained with anti-UNC-89 (Figure 5E). To characterize the depth of HA-UNC-68 localization at the muscle cell boundary, we co-stained HA-UNC-68 and PAT-6. As can be seen at the bottom of Figure 5F which present part of one muscle cell boundary, HA-UNC-68 is located more deeply than PAT-6.

**Figure 5.**
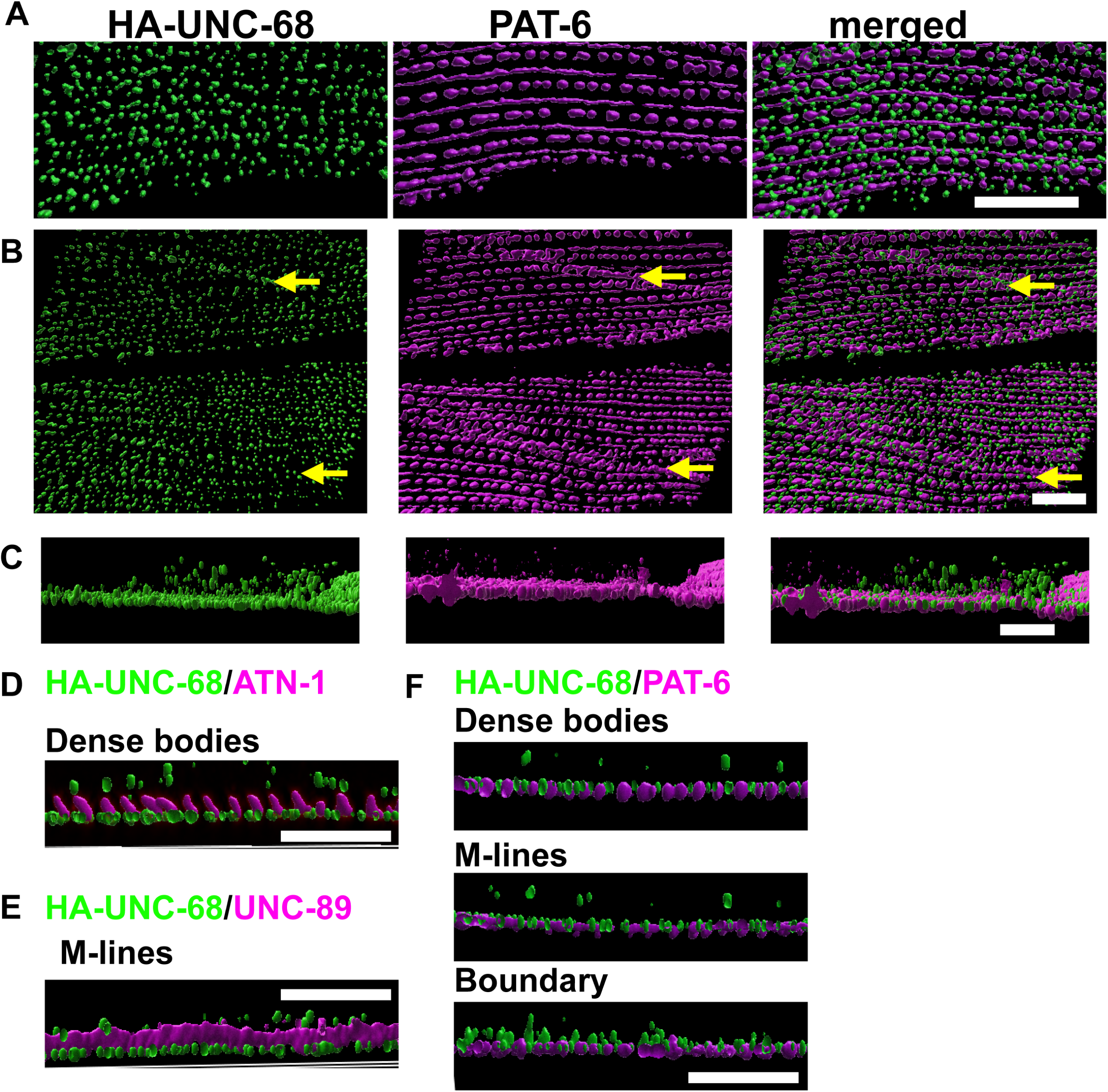
Localization of UNC-68 by structured illumination microscopy (SIM). (A-C) Portions of a single or multiple body wall muscle cells from strain SU980 (expresses HA-UNC-68) immunostained with antibodies to PAT-6 and antibodies to HA to visualize HA-UNC-68. PAT-6 is known to localize to the bases of M-lines and dense bodies, and to the adhesion plaques of muscle cell boundaries. (A) HA-UNC-68 localizes to puncta situated on either side of M-lines, and surrounding dense bodies, with the largest puncta being closest to the dense bodies. (B) With a wider field of view, HA-UNC-68 can be seen to also localize to muscle cell boundaries, as indicated by arrows. To create images shown in parts C-F, a Z-series of HA-UNC-68 and PAT-6 (or ATN-1 or UNC-89) images from the outer muscle cell membrane to deep into the muscle cell was used to create a 3D reconstruction. (C) Side view of 3D reconstruction of an entire muscle cell showing HA-UNC-68 and PAT-6 localization. For each panel, bottom is close to the outer muscle cell membrane. In addition to being localized like PAT-6 close to the muscle cell membrane, there is substantial localization of HA-UNC-68 in the deeper portions of the muscle cell. (D) Side view of part of a single row of dense bodies co-stained with HA-UNC-68 and ATN-1. ATN-1 is known to localize to the deepest portions of the dense bodies and defines the deepest portion of the myofilament lattice. HA-UNC-68 is found mainly near the outer muscle cell membrane but also extends deeper into the muscle cell than the myofilament lattice. (E) Side view of part of a single M-line co-stained with HA-UNC-68 and UNC-89. UNC-89 is known to localize to the entire depth of the M-line and also defines the extent of the myofilament lattice. HA-UNC-68 can be seen localized close to the muscle cell membrane and deeper into the cell than the myofilament lattice. (F) Side view of part of a single row of dense bodies, part of one M-line and part of one muscle cell boundary. Confirming the results in D and E, some HA-UNC-68 is found deeper than the myofilament lattice. In addition, at the muscle cell boundary HA-UNC-68 is found deeper than PAT-6. Scale bars, 5 μm.

Unfortunately, the anti-FARL-11 antibodies did not give a strong enough signal in immunostaining to allow SIM imaging. However, we were able to use confocal microscopy to acquire a Z series using anti-FARL-11 and anti-HA for HA-UNC-68, starting from the outer muscle cell membrane, to deeper into muscle cells (Figure 6). FARL-11 is found between and around dense bodies throughout their depth and is also found at muscle cell boundaries (indicated by arrows). Curiously, HA-UNC-68 only appears at the muscle cell boundaries at the deepest portions of the muscle cell, again consistent with the SIM images. Localization of UNC-68 at the muscle cell boundaries has not previously been reported and its possible function there is unknown. The function of FARL-11 at muscle cell boundaries, particularly throughout the depth of the boundaries of muscle cells, is also unclear. Overall, FARL-11 localization is similar to that of HA-UNC-68 in Z-series, suggesting that FARL-11 is localized to the SR.

**Figure 6.**
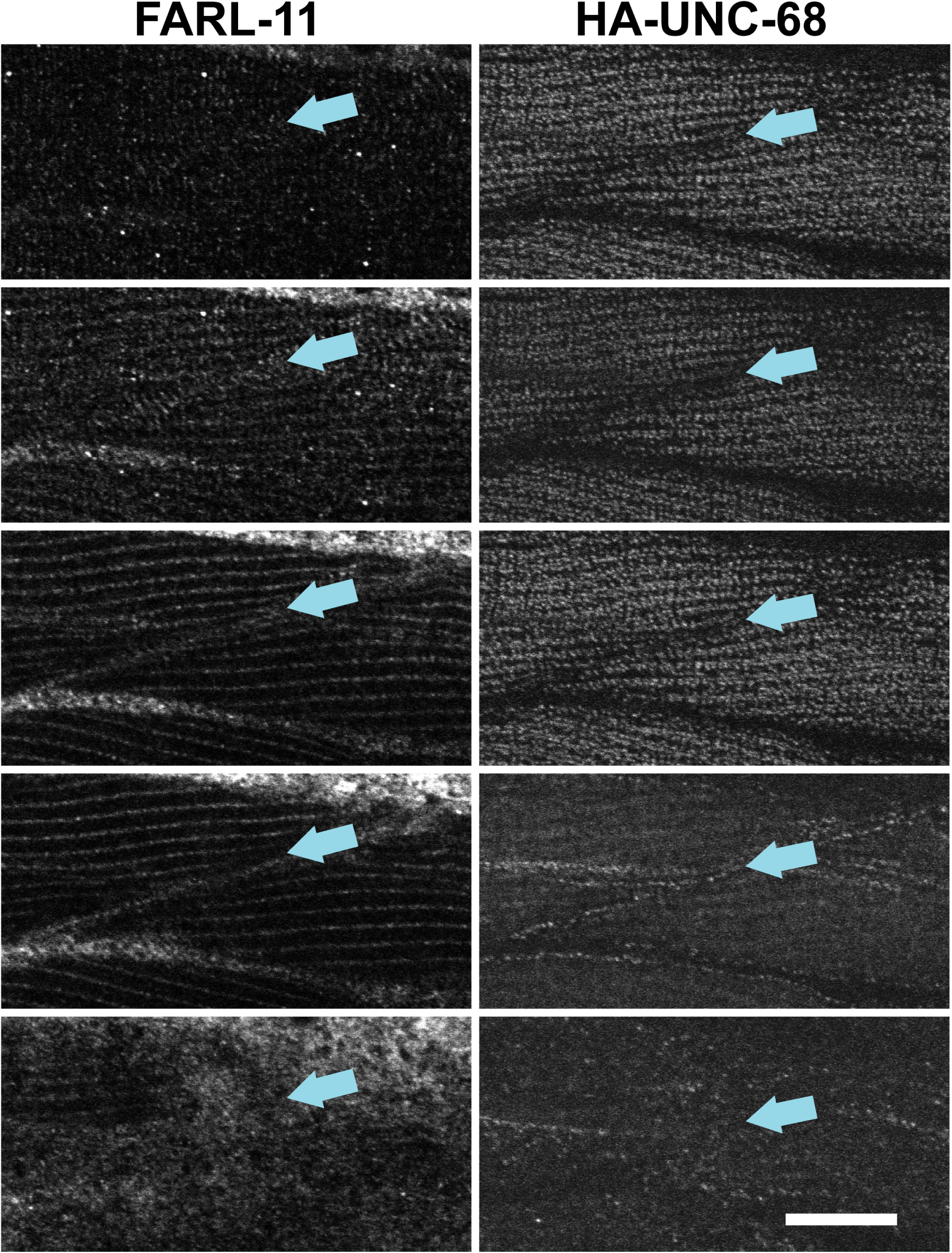
A confocal Z-series reveals that FARL-11 localization is similar to localization of HA-UNC-68. Strain SU980 was co-stained with antibodies to FARL-11 and to HA. From top to bottom are 0.5 μm-thick sections from the outer portion of a body wall muscle cell consecutively deeper into the muscle cell. FARL-11 is found between and around dense bodies throughout their depth and is also found at muscle cell boundaries (indicated by the arrows). HA-UNC-68 only appears at the muscle cell boundaries at the deepest portions of the muscle cells (2^nd^ row from the bottom of the figure). Scale bar, 10 μm.

In mammalian muscle, the UNC-89 homolog, obscurin, links myofibrils to the surrounding SR, through interaction of obscurin with the SR membrane proteins small ankyrin 1 and 2 (Bagnato et al., 2003; Kontrogianni-Konstantopoulos et al., 2003). Moreover, knockout of the mouse obscurin gene results in disorganization of the SR (Lange et al., 2009). This role for obscurin/UNC-89 seems to be evolutionarily conserved. In *C. elegans*, there is genetic evidence for an UNC-89 to SR functional linkage. VAV-1 is a RacGEF that regulates the concentration of intracellular calcium and is expressed in body wall muscle; overexpression of *vav-1* in muscle results in slow movement; mutagenesis resulted in suppressors that move better; the suppressor mutations are in *egl-19* (an L-type calcium channel) and *unc-89* (Spooner et al., 2012). Moreover, in *unc-89* mutants, in addition to disorganization of the sarcomeres, there is disorganization of the SR as probed using transgenics overexpressing MYC-UNC-68 or SERCA-GFP (Spooner et al., 2012). Therefore, we wondered whether loss of function of *unc-68*, which has been shown to disrupt SR organization (Maryon et al., 1998) might also disrupt the organization of sarcomeres. As shown in Figure 7, *unc-68(e540)* shows disorganization of major structural components of the sarcomere—the A-bands (MHC A), the M-lines (UNC-89), and the bases of M-lines and dense bodies (UNC-95).

**Figure 7.**
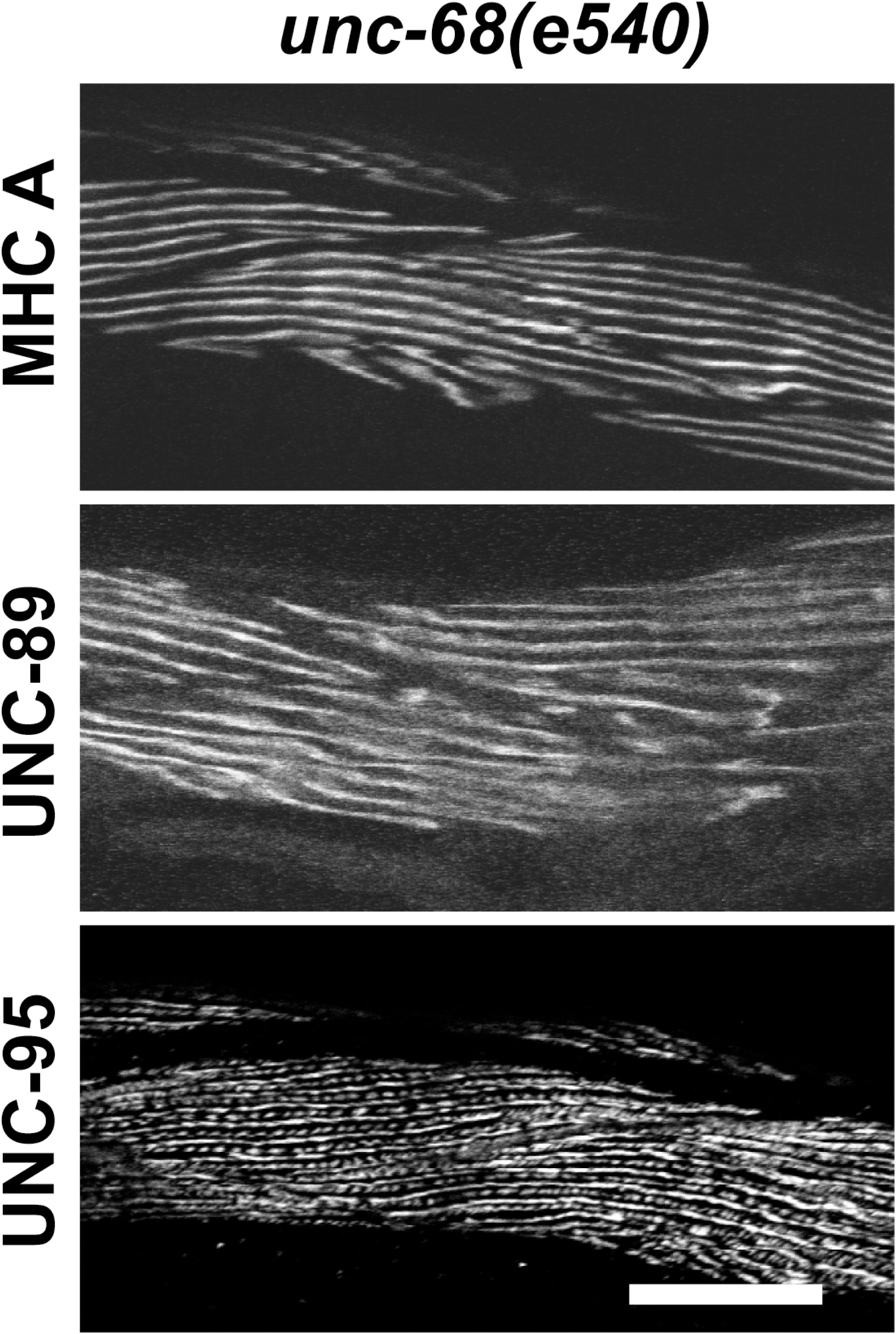
Loss of function of *unc-68*, which encodes the SR calcium release channel, results in disorganized sarcomeres. The loss of function mutant *unc-68(e540)* was immunostained with antibodies to MHC A myosin, UNC-89, and UNC-95. Each row shows representative images from two different animals. All the structures labeled by these antibodies (A-bands, M-lines, dense bodies and muscle cell boundaries) show significant disorganization (compare to wild type shown in Figure 2B). Scale bar, 10 μm.

If, as our data suggests, FARL-11 and CASH-1 are components of the SR, one straightforward prediction is that in a mutant in which the SR is disrupted, e.g. an *unc-68* mutant, FARL-11 and CASH-1 might be mislocalized. Curiously, however, this is not the case. As shown in Figure 8A, the localization of FARL-11 in *unc-68(e540)* and wildtype are nearly identical. Because *cash-1* and *unc-68* are very close together on the genetic map, we used CRISPR to tag CASH-1 at the N terminus using TagRFP::3xmyc in an *unc-68(e540)* mutant background. As shown in Figure 8B, the localization of CASH-1 in *unc-68(e540)* and wildtype are nearly identical.

**Figure 8.**
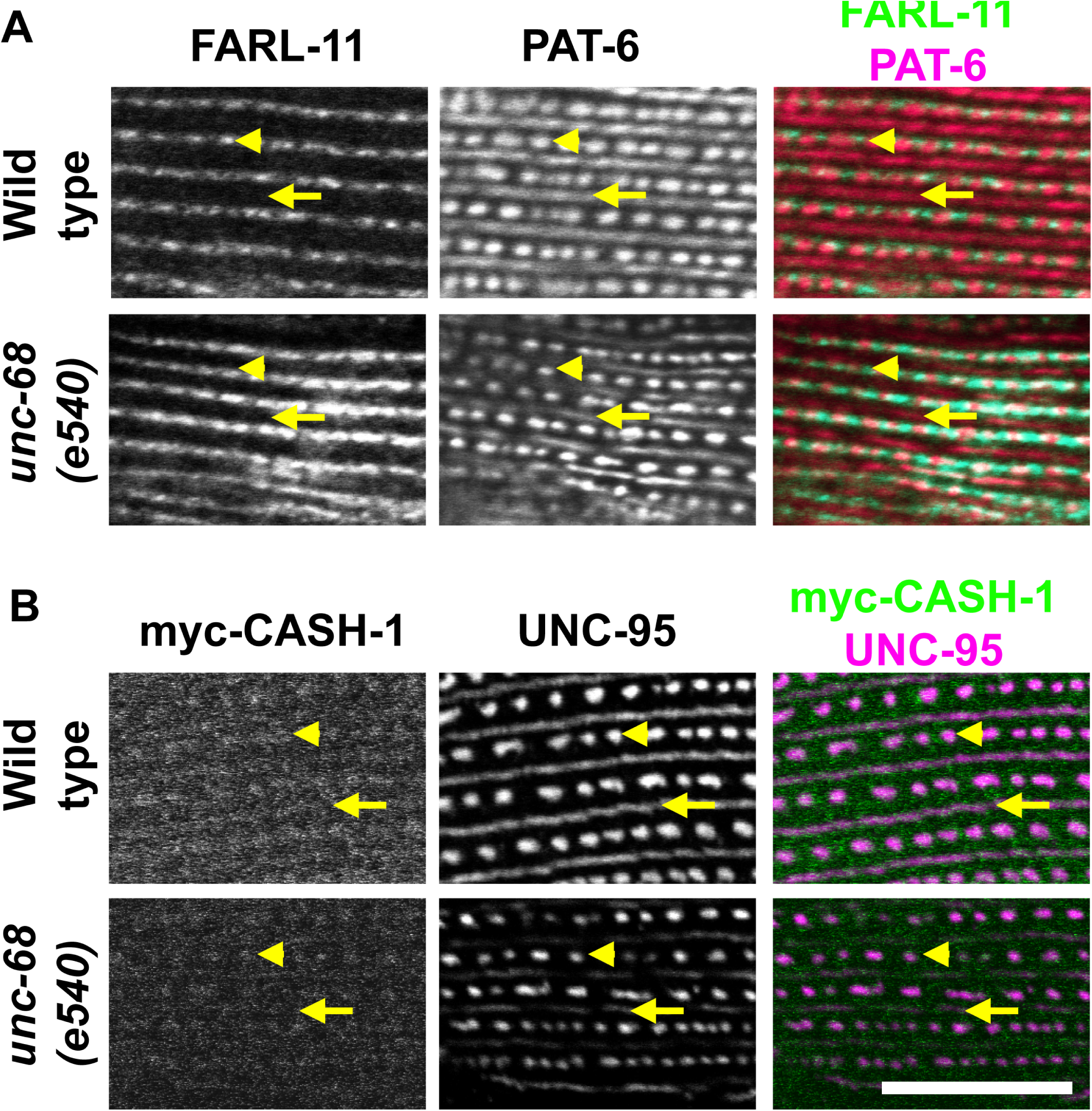
Loss of function of *unc-68* does not affect the localization of FARL-11 or CASH-1. (A) Wild type and *unc-68(e540)* were co-stained with antibodies to FARL-11 and PAT-6. Note that the localization of FARL-11 in *unc-68(e540)* is no different from the localization of FARL-11 in wild type. (B) SU854 which expresses RFP and myc tagged CASH-1, designated “wild type”, and SU1086 which expresses RFP and myc tagged CASH-1 and also has the *unc-68(e540)* mutation, designated “*unc-68(e540)”* were co-stained with antibodies to myc to detect myc-CASH-1 and antibodies to UNC-95 to mark the bases of M-lines and dense bodies. As shown, there is no obvious difference in the localization of myc-CASH-1 in wild type vs. *unc-68(e540).* Scale bar, 10 μm.

We next tested the hypothesis that FARL-11 and CASH-1 proteins affect the organization of the SR or at least the SR marker UNC-68. To address this, we created two strains, one in which HA-UNC-68 is expressed in *farl-11(gk437008),* and one in which HA-UNC-68 is expressed in *cash-1(jc100)*. As compared to the localization of HA-UNC-68 in a wild type background (Figure 9A), the overall density of puncta is reduced, especially near the M-lines, in a *farl-11(gk437008)* mutant background (Figure 9B). However, in a *cash-1(jc100)* mutant background, we observed no change in overall pattern of HA-UNC-68 puncta (Figure 9C)

**Figure 9.**
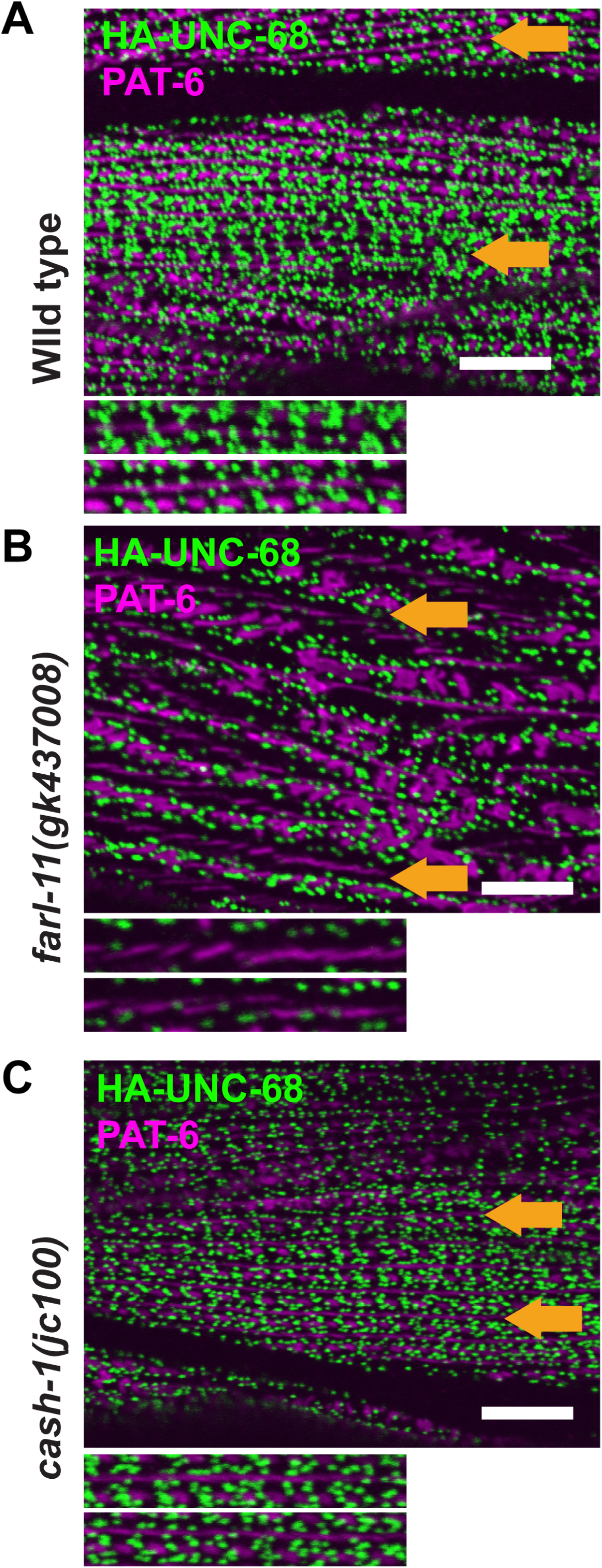
The organization of UNC-68 is adversely affected by loss of function of *farl-11*. Confocal imaging of body wall muscle co-stained with antibodies to HA to detect HA-UNC-68, and antibodies to PAT-6. Scale bar, 10 μm. (A) Imaging of SU980 which expresses HA-UNC-68 in a wild type background. The two thin rectangles at the bottom are zoomed-in images of the areas denoted by the orange arrows. Note the accumulation of HA-UNC-68 small puncta around M-lines (magenta lines of PAT-6 staining). (B) Imaging of GB352 which expresses HA-UNC-68 in a *farl-11(gk437008)* mutant background. Note that in the zoomed-in images that there are fewer HA-UNC-68 small puncta surrounding M-lines, as compared to what is observed in a wild type background shown in (A). (C) Imaging of SU1070 which expresses HA-UNC-68 in a *cash-1(jc100)* mutant background. Note that in the zoomed-in images that UNC-68 small puncta have the same density and arrangement of distribution around M-lines as in the wild type background shown in (A).

Given that the overall density of HA-UNC-68 puncta is reduced in the *farl-11* mutant, we wondered if there would also be a decrease in the overall level of HA-UNC-68 protein. To address this question, we performed a western blot in which protein extracts from wild type, *farl-11(gk437008)* and *cash-1(jc100)* were separated on a 5% polyacrylamide gel. As shown in Figure 10, anti-HA detects two major protein bands, one running at the expected size of 557-572 kDa (indicated by a blue arrow), and the other running at ∼180 kDa (indicated by a yellow arrow). The origin of the ∼180 kDa protein is unknown but we suspect that it is results from either normal processing of UNC-68, or from degradation during extract preparation. Quantitation from three experiments shows that, as compared to wild type, (1) the level of the 560 kDa band is not changed in *farl-11*, but is increased in *cash-1*; (2) the level of the 180 kDa band is reduced in both *farl-11* and *cash-1*; and (3) the combined levels of both the 560 and 180 kDa bands are reduced in both *farl-11* and *cash-1*. Thus, we can conclude that the status of *farl-11* and *cash-1* affects the level of UNC-68, consistent with the reduced number of puncta observed in the *farl-11* mutant.

**Figure 10.**
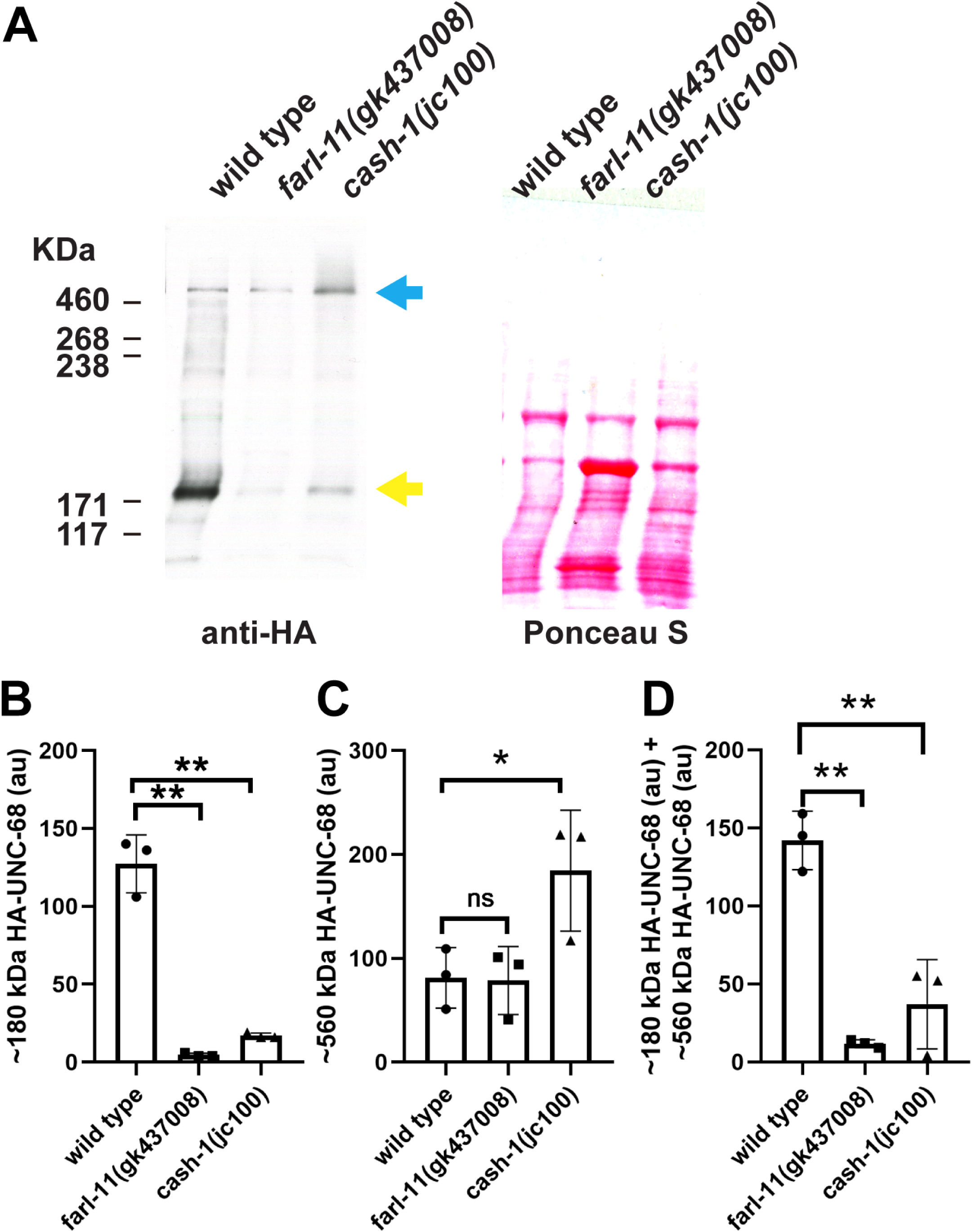
The level of HA-UNC-68 and/or its fragments are affected by loss of function of *farl-11* and *cash-1*. (A) Total protein extracts from strains expressing HA-UNC-68 in wild type, *farl-11* and *cash-1* mutant backgrounds were separated on 5% SDS-PAGE, transferred to membrane and reacted with antibodies to HA. On the left is the western reaction, and on the right is the same membrane stained with Ponceau S. Molecular weight size markers are shown on the far left. The arrow indicates the position of the full-length HA-UNC-68, which is expected to be 557-572 KDa (multiple isoforms); the asterisk represents a likely N-terminal ∼180 KDa fragment from HA-UNC-68. (B,C,D) Quantitation of the levels of the 180 kDa, the 560 kDA and the sum of both proteins, comparing wild type vs. f*arl-11* and *cash-1* mutants. For each strain N=3, and significance was tested by a one-tailed student’s t test.

## Discussion

From this study, we have expanded the function of PP2A in striated muscle. We previously reported that loss of function of the catalytic subunit (LET-92) or the scaffolding subunit (PAA-1) results in muscle cells detaching from the ECM, and that loss of function of various regulatory (B) subunits results in disorganization of sarcomeres. We had also demonstrated that various components of the PP2A complex localize to various sarcomeric structures (I-band, M-lines, and dense bodies (Z-disks)). We had shown that RNAi for regulatory B’’’ subunit CASH-1 (STRN3) results in disorganized sarcomeres, however we did not have reagents available to localize the CASH-1 protein. In the current study, we took into account that striatin forms a special PP2A complex, associating not only with catalytic and scaffolding subunits of PP2A but also with STRIP1. The *C. elegans* ortholog of STRIP1 is called FARL-11, and FARL-11 localizes to the ER and the outer nuclear membrane of early embryos and is required for normal ER organization. This prompted us to determine if FARL-11 has a role in the muscle specific ER, the sarcoplasmic reticulum (SR). Consistent with the known structure of the human STRIPAK complex, we found that CASH-1 (STRN3) and FARL-11(STRIP1) form a complex in vivo (Figure 1C).

Similar to loss of function of canonical PP2A subunits (Qadota et al., 2018), loss of function of CASH-1 (STRN3), or FARL-11 (STRIP1), show disorganized sarcomeres, including a variable broadening of the distribution of myosin and especially UNC-89 (obscurin) (Figures 2 and 3). We found that both CASH-1 and FARL-11 localize between and surrounding dense bodies (Figure 4). We used CRISPR to add an HA tag to the N-terminus of UNC-68 (ryanodine receptor), and localized HA-UNC-68 by SIM to a series of puncta located on either side of the M-lines, and surrounding dense bodies, with the larger puncta being closer to the dense bodies (Figures 5 and 9). Co-staining of anti-HA for HA-UNC-68 together with antibodies to FARL-11, or with antibodies to myc for myc-CASH-1, and conventional confocal microscopy, reveals that HA-UNC-68 and myc-CASH-1 partially co-localize (Figure 4 D and E, Figure 6). Unfortunately, neither the anti-FARL-11 antibodies nor the anti-myc antibodies yielded a strong enough signal to allow SIM imaging. However, a Z series acquired by confocal microscopy after staining with anti-FARL-11 and anti-HA for HA-UNC-68, shows that FARL-11 localization is similar to that of HA-UNC-68 (Figure 6), consistent with FARL-11 being localized to the SR. If CASH-1 and FARL-11 are required for the formation or maintenance of SR, we would expect that disorganization of SR might not affect the localization of CASH-1 or FARL-11, but loss of CASH-1 or FARL-11 might affect the organization of the SR. Our results were consistent with this expectation: an *unc-68* mutant does not affect the localization of FARL-11 or CASH-1 (Figure 8); however, a *farl-11* mutant shows overall reduced density of UNC-68 puncta, especially near the M-lines (Figure 9). Moreover, by western blot, the overall level of UNC-68 is reduced in both *farl-11* and *cash-1* mutants as compared to wild type (Figure 10).

An interesting effect of mutations in *farl-11* or *cash-1* is a widening of the localization of UNC-89 and MHC A myosin (Figure 2B, Figure 3C). The fact that both UNC-89 and MHC A are affected can be explained by considering that UNC-89 is located throughout the depth of the M-line (Warner et al., 2013), the M-line is the region of the sarcomeric A-band where thick filaments are cross-linked, and MHC A is the myosin isoform that is restricted to the middle of the thick filament in *C. elegans* (Miller et al., 1983). But how loss of function of *farl-11* or *cash-1* results in widening of the M-line region components is more difficult to explain. One possibility is suggested by our finding that there are reduced numbers of UNC-68-containing puncta around the M-lines (Figure 9). Perhaps the SR sacs that lie on either side of the base of the M-line, restrict the growth of the M-line laterally, either by physical hindrance or signaling. In support of this idea, in mammalian striated muscle, obscurin at the M-line links sarcomeres to the surrounding SR by interacting with SR membrane proteins small ankyrins 1 and 2 (Bagnato et al., 2003; Kontrogianini-Konstantopoulos et al., 2003). Another possibility is that with the reduced amount of SR in the *farl-11* mutant, Ca^+2^ signaling is disturbed and this results in unregulated or asymmetric contraction which ultimately results in disorganization of the sarcomeres.

We also found that FARL-11, and for the first time that UNC-68, are located at the muscle cell boundaries (Figures 5 and 6), in addition to their locations surrounding M-lines and dense bodies. In addition, a Z-series indicates that UNC-68 only appears at the muscle cell boundaries at the deepest portions of the muscle cell. It should be noted that *C. elegans* spindle-shaped body wall muscle cells are arranged cell to ECM to cell, where attachment plaques of each cell anchor the cell to a thin layer of ECM that lies between the adjacent cells. These attachment plaques are integrin adhesion complexes and contain a subset of proteins found at dense bodies (Qadota et al., 2017). Electron microscopy reveals that the muscle boundaries also contain gap junctions and finger-like projections of one cell into another cell (Qadota et al., 2017). We had previously speculated that the gap junctions and finger-like projections provide electrical conductivity, signaling and structural integrity allowing multiple smaller cells to function as larger units. Although membranous sacs reminiscent of SR have not been found at muscle cell boundaries, perhaps UNC-68 embedded in the cell membranes might facilitate movement of Ca^+2^ between muscle cells.

Finally, we would like to point out that we took advantage of the Million Mutation Project to identify and study missense mutants in conserved residues in *cash-1* and *farl-11* that developed into adults and were fertile. This allowed us to more easily study adult muscle phenotypes. For example, an intragenic deletion of *farl-11*, *tm6233*, is noted on WormBase to be lethal or sterile.

## Methods

### *C. elegans* Strains

N2 (wild type, Bristol)

SU853 *farl-11 (jc61[mNG::TEV::3xflag::farl-11 + LoxP])* III

SU854 *cash-1 (jc60[TagRFP::TEV::3xmyc::cash-1 +Lox 2272])* V

SU980 *unc-68*(jc78[*3xha::unc-68*]) V

SU1002 *cash-1* (*jc60*[*[TagRFP::TEV::3xmyc::cash-1 +Lox 2272*]; *unc-*

*68*(jc78[*3xha::unc-68*]) V-outcrossed to wild type 2X (o.c. 2x)

SU1047 *farl-11* (j*c93* [*farl-11Δ T281 *jc61*]) II -o.c. 2x

SU1048 *farl-11* (*jc94* [*farl-11 ΔP280 *jc61*]) II

SU1054: *cash-1* (*jc100*[*cash-1 h121y *jc60*]) V-o.c. 2x

SU1086 *cash-1* (*jc106[3xmyc::cash-1*]); *unc-68* (*e540*)-o.c. 2x

SU1070 *cash-1* (*jc100*[*cash-1h121y *jc60*]); *unc-68* (*jc78* [*3xha::unc-68*a])-V-o.c. 2x

CB540 *unc-68 (e540)* V

VC40583 *cash-1* (*gk705529*[*cash-1 H121Y*])-V

VC30204 *farl-11* (*gk437008* [*farl-11 L292P*])-II

GB351 *farl-11(gk437008)* [FARL-11 L292P]-II o.c. 3x

GB352 *farl-11(gk437008)* [FARL-11 L292P]-II o.c. 3x; *unc-68* (*jc78* [*3xha::unc-68*a])-V

SU1030 cash-1(syb4646[TagRFP::CASH-1 L82E/L86E*jc60])V / tmC12[egl-

9(tmIs1197)] V

PHX4646 *cash-1*(*syb4646* [*TagRFP::cash-1 L82E/L86E *jc60*])V / *nT1* (*qIS51*)] (IV; V)

### CRISPR/Cas9

To generate endogenous insertions a self-excising cassette was used as described (Dickinson *et al.,* 2015). *tagrfp::SEC::CASH-1/+* was crossed into the *tmC12 (*Dejima *et al.,* 2018) balancer to facilitate SEC removal. *mng::SEC::farl-11/+* was crossed into the *mIn1* balancer to facilitate SEC removal. The balanced progeny were heat shocked at 34°C to remove the SEC. SEC excision and correct knock-in were confirmed by PCR of non-rolling, non-balanced progeny. Worms were outcrossed with N2 a minimum of 2x.

Sunybiotech generated the PHX4646 *cash-1*(*syb4646* [*TagRFP::cash-1 L82E/L86E *jc60*])V / *nT1* (*qIS51*)] (IV; V) strain. This strain was outcrossed with N2 and then crossed into the *tmC12* [*egl-9*(*tmIs*1197)] strain to create SU1030 *cash-1* (*syb4646* [*TagRFP::cash-1 L82E/L86E *jc60*])V/*tmC12* [*egl-9*(*tmIs*1197)] V

Single point mutants were generated as described (Arribere *et al.,* 2014).

Guide RNA sequences were cloned into the Cas9 containing plasmid (pJW1219).

CASH-1 TagRFP 5’ guide- 5’ -CGGATTCGAGTAGTTACAATGG

CASH-1 H121Y guide- 5’ ATTTATGTATGCATCAAGGT

CASH-1 L28E/L86E guide 1- GCTCGTATTGCTTTCCTACAAGG

CASH-1 L82E/L86E guide 2 - AATTATCGACTAACCCACAACGG

FARL-11-mNG 5’ guide: - 5’GGCCTGATTCCCATTACATT

FARL-11 L290P guide- 5’ ATTTCATTATCAACCATAGT

UNC-68-5’guide-5’ TCCTCCCTGCTCCTCCTTGT

### Homology searches/protein sequence analysis

CASH-1 and FARL-11 protein sequences were run through a Hydrophobic Cluster analysis program (mboyle@RPBS) to generate a Hydrophobic Cluster Plot (Néron *et al.,* 2009*). jc93*, *jc94*, and *jc100* were generated at sites of predicted secondary structure (Callebaut *et al*., 1997). *cash-1 L82E* /*L86E* mutations were designed to mimic the mutations in the homodimerization domain of STRN-3 (Chen *et al.,* 2014)

### Protein structure modeling

For FARL-11 (uniprot Q19300) and CASH-1 (uniprot G5EE12) protein structure modeling, CLUSTALW version 1.2.2 (https://www.ebi.ac.uk/Tools/msa/clustalw2/), SWISS-MODEL version July 2021 (https://swissmodel.expasy.org/; Waterhouse et al., 2018) and Phyre2 version 2.0 (http://www.sbg.bio.ic.ac.uk.phyre2/html/page.cgi?id=index; Kelley LA et al., 2015) online tools were used.

For FARL-11, we used the cryoEM structure of human ortholog of STRIP1 as reference for modeling (7K36.pdb; Jeong et al., 2021). For CASH-1, we used the human ortholog STRN3 as reference for modeling (7K36.pdb; Jeong et al, 2021). Molecular graphics were generated by using Chimera version 1.15 https://www.cgl.ucsf.edu/chimera/; Pettersen et al., 2004). Single amino acid mutations were inserted using the rotamer tool and then energy minimized to minimize interatomic clashes and contacts based on van der Waals radii. In addition, we used the DynaMut2 (Rodrigues et al.. 2021) to predict the effects of the point mutations on protein stability and dynamics.

### FARL-11 antibody construction

Antibody production was performed by Li International (Denver, CO). A 50 amino acid chemically synthesized peptide corresponding to the extreme C-terminus of FARL-11 (aa 929-978

TCAHSVLGANLKLGRHFKKDYEKWLEQEVFNASIDWDKLLIETRGVEDLM) was injected into New Zealand rabbits. Li International performed immunization and provided lyophilized antibody which was resuspended to 1 mg/mL with 1x PBS.

### Co-immunoprecipitation of FARL-11 with CASH-1

Co-immunoprecipitation was performed essentially as described (Zaidel-Bar et al. 2010). 20, 10 cm NGM plates with gravid adults were harvested and frozen in Lysis (-) buffer (50 mM HEPES pH 7.4, 150 mM NaCl, 0.05 mM DTT). Worms were lysed by sonication in Lysis (+) buffer (50 mM HEPES pH 7.4, 150 mM NaCl, 0.05 mM DTT, 1% NP-40) and total lysates were centrifuged at 4° C. The supernatant was then pre-cleared using Chromotek control magnetic agarose beads (Chromotek, cat. no. gmab-20) for 30 minutes at 4° C. Pre-cleared lysates were then incubated with Chromotek RFP-Trap magnetic agarose beads (cat. no. rtma-20) for 1 hour at 4° C to bind to TagRFP-myc-CASH-1. Beads were washed 3X with 1 ml of Lysis (-) buffer, and eluted with 50 ml of Lysis (-) buffer at 90° C for 10 minutes. Samples were resolved on 10% SDS-PAGE and transferred to PVDF membrane (Immobilon-FL, Millipore Corp., cat. no. IPFL00010) using a ThermoFisher Powerblotter (cat. no. PB0013), and immunoblotting was performed using ThermoFisher iBindFlex (cat. no. SLF2000). FARL-11 was detected at 1:500 dilution using the rabbit anti-FARL-11 antibody described above. LiCor 2° goat anti-rabbit antibody (cat. no. 925-3211) was used for detection at 1:4,000 dilution.

### Western blots

The method of Hannak et al. (2002) was used to prepare total protein lysates from mixed stage populations of wild-type, SU853, SU854, SU1054, *farl-11(jc93)*, *farl-11(jc94)*, *farl-11(gk437008)*, SU980, GB352 and SU1070 worms. For the western blots presented in Figures 1-3, equal amounts of total protein were separated on 10% polyacrylamide-SDS-Laemmli gels, transferred to nitrocellulose membranes, reacted with affinity purified, anti-FARL-11 at 1:500 or 1:1000 dilution, and anti-myc (mouse monoclonal clone 9E 10 from the University of Iowa Hybridoma Bank) at 1:300 dilution. Blots were reacted with anti-rabbit (or anti-mouse) immunoglobulin G conjugated to HRP (GE Healthcare) at 1:10,000 dilution, and visualized by ECL. For the western blot comparing the level of HA-UNC-68 in wild type vs. farl-11 an cash-1 mutant backgrounds shown in Figure 10, equal amounts of total protein from each of these strains were separated by a 5% separating and 3% stacking Laemmli SDS-PAGE gel, transferred to nitrocellulose membrane for 2 h, and reacted with anti-HA (rabbit monoclonal antibody from Cell Signaling Technologies, cat. no. 3724S) at 1:1,000 dilution, followed by reaction with anti-rabbit immunoglobulin G conjugated to HRP (GE Healthcare) at 1:10,000 dilution and visualized by ECL. For protein molecular weight size markers, we used the HiMark Pre-Stained Protein Standard (cat. no. LC5699 from Life Technologies). For comparison of protein levels between wildtype and mutants, samples were normalized based on total protein per lane visualized by Ponceau S staining.

### Immunostaining, confocal and SIM microscopy of body wall muscle

Adult nematodes were fixed and immunostained using the method described by Nonet et al. (1993) with further details described in Wilson et al. (2012). The following primary antibodies were used at 1:200 dilution except as noted: anti-UNC-89 (mouse monoclonal MH42; Benian et al., 1996; Hresko et al., 1994), anti-MHC A (mouse monoclonal 5-6; Miller et al., 1983), anti-UNC-95 (rabbit polyclonal Benian-13; Qadota et al., 2007), anti-PAT-6 (rat polyclonal; Warner et al., 2013), anti-ATN-1 (mouse monoclonal MH35; Francis and Waterston, 1991), anti-HA (rabbit monoclonal from Cell Signaling Technology Inc., cat. no. 3724S; and mouse monoclonal from Sigma-Aldrich, cat. no. H3663), anti-myc (mouse monoclonal clone 9E 10 from the University of Iowa Hybridoma Bank), and anti-FARL-11 (this study). Secondary antibodies, used at 1:200 dilution, included anti-rabbit Alexa 488, anti-rat Alexa 594, and anti-mouse Alexa 594, all purchased from Invitrogen. Images were captured at room temperature with a Zeiss confocal system (LSM510) equipped with an Axiovert 100M microscope and an Apochromat x63/1.4 numerical aperture oil immersion objective, in 1x and 2.5x zoom mode. Super-resolution microscopy was conducted with a Nikon N-SIM system in 3D structured illumination mode on an Eclipse Ti-E microscope equipped with a 100×/1.49 NA oil immersion objective, 488- and 561-nm solid-state lasers, and an EM-CCD camera (DU-897, Andor Technology). Super-resolution images were reconstructed using the N-SIM module in NIS-Elements software. For all images, confocal and SIM, color balance was adjusted by using Adobe Photoshop (Adobe, San Jose, CA).

## Acknowledgements

H.Q. and G.M.B. were supported by NIH grant R01HL160693 and NSF grant 2050009. J.H. and S.C.T.M. were supported by NIH grants R01GM058038 and MIRA R35GM145312. A.F.O. acknowledges support from The Cecil and Ida Green Endowment. In addition, S.C.T.M. was supported by a NIH T32 (GM130550) Biophysics Training Grant, a Gilliam Fellowship from the Howard Hughes Medical Institute, an Advanced Opportunities Fellowship, and a COVID-19 dissertation completion fellowship from the University of Wisconsin-Madison.

## Author contributions

S.C.T.M. and H.Q. designed, carried out and interpreted most of the experiments and wrote some sections of the manuscript; A.F.O. created the protein homology models and interpreted the mutations; J.H. and G.M.B. conceived of the project, provided funding and edited the manuscript; G.M.B. carried out a few experiments and wrote most of the paper.

---

**Supplemental Figure 1.** Imaging of GB352 which expresses HA-UNC-68 in a *farl-11(gk437008)* mutant background, immunostained with anti-HA and anti-UNC-89, an alternative way to mark M-lines (shown in magenta). Note that in the zoomed-in images that there are fewer HA-UNC-68 small puncta surrounding M-lines, as compared to wild type shown in Figure 9A. Scale bar, 10 μm.

