## Supplementary figures and images for "FARL-11 (STRIP1/2) is Required for Sarcomere and Sarcoplasmic Reticulum Organization in *C. elegans*"

### Supplemental Figure 1

#### Supplemental Figure 1

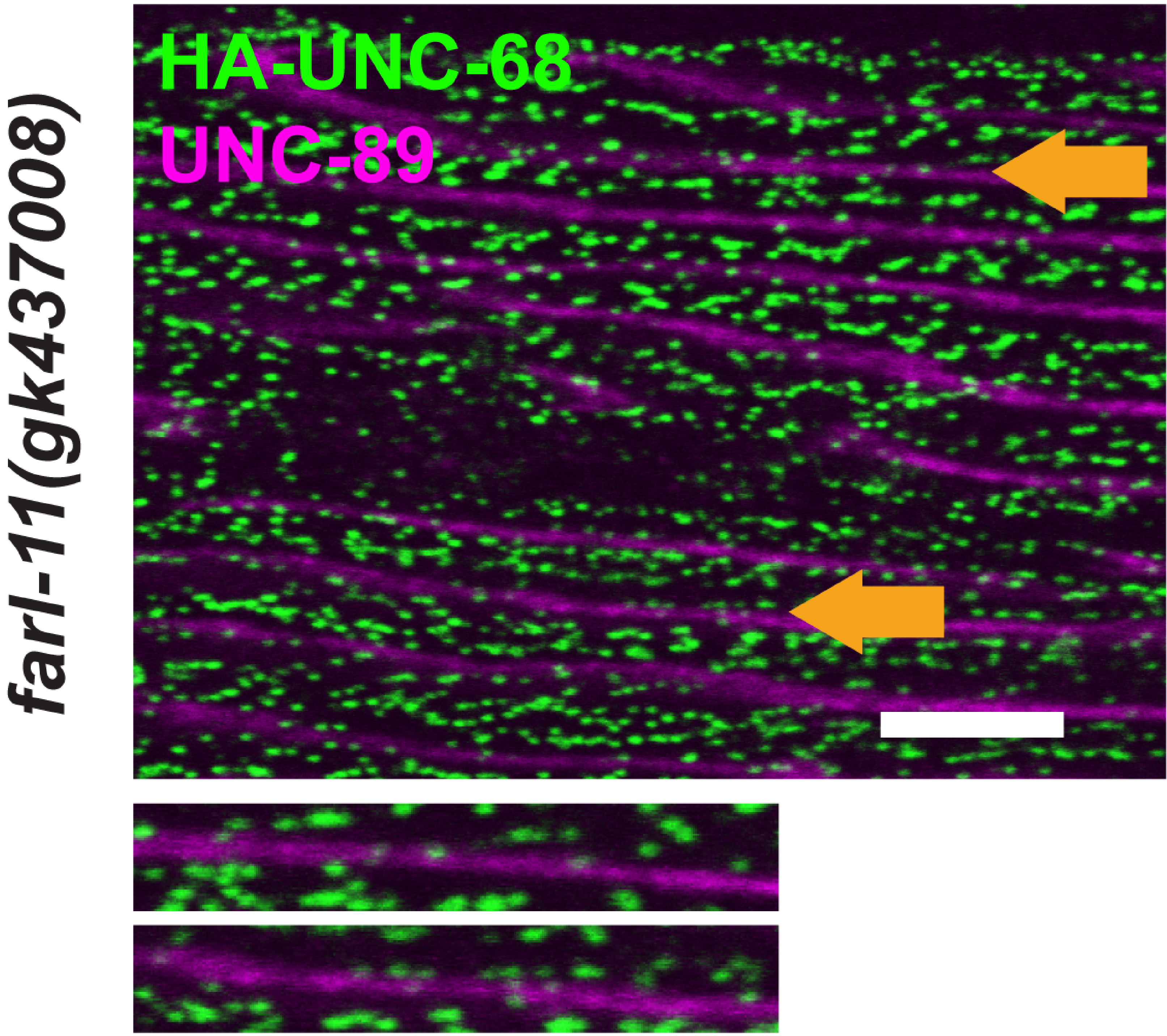
